# *Rickettsia japonica* in ticks infesting a wild mammal in Thailand

**DOI:** 10.1101/2021.02.23.432398

**Authors:** Supanee Hirunkanokpun, Arunee Ahantarig, Visut Baimai, Pairot Pramual, Wachareeporn Trinachartvanit

**Affiliations:** Ramkhamhaeng University, Bangkok, Thailand; Mahidol University, Phayathai, Bangkok, Thailand; Mahasarakham University, Maha Sarakham, Thailand

**Author notes:** **Address for correspondence**: Wachareeporn Trinachartvanit, 272 Department of Biology, Faculty of Science, Mahidol University, Rama 6 Road, Ratchathewi, Bangkok 10400, Thailand.

**Keywords:** *Rickettsia japonica*, Spotted fever group, ticks, Thailand

## Abstract

We report the first case of *Rickettsia japonica* in *Haemaphysalis hystricis* ticks collected from a Burmese ferret-badger in Loei province, northeastern Thailand. Phylogenetic analyses of *omp*A and *omp*B genes clearly showed that it was almost identical to *R. japonica* found in ticks and Japanese spotted fever (JSF) patients previously reported.

## Text

Ticks (Acari: Ixodidae) have been considered as important vectors of various infectious agents in Southeast Asia including spotted fever group (SFG) rickettsiae, and especially *Rickettsia japonica* causing Japanese spotted fever (JSF) which can be transmitted to humans through infected ticks (*1*). JSF has mainly been reported in Japan, South Korea, China, and the Philippines (*1*). In Thailand, human infection with *Rickettsia* sp. related to *R. japonica* was first reported in a male patient (*2*). Later, the strain TCM1 of *Rickettsia* sp. related to *R. japonica* was successfully isolated from a *Haemaphysalis hystricis* tick found on Mt. Doi Suthep, northern, Thailand, but the identity of its host was not reported (*3*). Although ticks are known as the potential vectors of SFG rickettsiae (*1,4*), in Thailand, there is very little known about interaction between vectors and mammalian host reservoirs.

To investigate the association of mammalian hosts parasitized by ticks infected with SFG rickettsiae, we examined 8 nymphs of *H. hystricis* removed from a fresh roadkill Burmese ferret-badger (*Melogale personata*) in northeastern Thailand centration in October 2019. Each individual nymph was washed in 10% bleach, 70% ethanol, and sterile distilled water three times (5 min each). DNA was extracted from 2 pooled nymphs (4 nymphs/pool) using the QIAamp DNA Extraction Kit for Tissue (QIAGEN) according to the manufacturer’s protocol. The presence of *Rickettsia* spp. was examined by Polymerase Chain Reaction (PCR). One pool of nymphs was PCR-positive and was subjected to DNA sequencing targeting the partial rickettsial 17-kDa antigen gene (*5*), the citrate synthase gene (*glt*A), outer membrane protein A (*omp*A) (*6*) and B (*omp*B) genes. In this study, a new set of primers was designed to amplify the rickettsial *omp*B region (800 bp product size); RicF: CAG CAA GGT AAT AAG TTT AAT AC and RicR: GCT ATA CCG CCT GTA GTA ACA G; Cycling conditions in PCR were 95°C for 5 min, 30 cycles of 95°C for 1 min, 56°C for 1 min, 72°C for 1 min, and a final cycle of 72°C for 10 min. The sequences for *Rickettsia* sp. in pooled nymphs of *H. hystricis* from Thailand (HHT) used in this study were submitted to the GenBank accession numbers MW415892 (17-kDa antigen), MW415894 (*glt*A), MW415896 (*omp*A), and MW415898 (*omp*B).

BLASTn analyses revealed that the rickettsial partial sequences of 17-kDa antigen and *glt*A genes were identical to several strains of *R. japonica, R. vini*, and *R. raoultii* deposited in GenBank. Since these two genes are highly conserved they are consequently not appropriate to differentiate species and/or strains of *Rickettsia* in this study. In contrast, the *omp*A and *omp*B genes are more variable and they therefore were employed in this study. BLAST results revealed that the *omp*A sequence obtained from *H. hystricis* nymphs removed from the Burmese ferret-badger was nearly identical (99.57%) to *R. japonica* infected humans in China (access. no. CP047359, KY347792-3), Japan (access. no. AP017581, LC101443, U43795), Korea (access. no. DQ019319), and the strain TCM1 isolated from *H. hystricis* in Thailand (access. no. AB359459). However, the sequence was quite distantly related to *Rickettsia* sp. infecting humans in Thailand (access. no. DQ909072) (Figure). The sequence of the *omp*B gene of the same sample was identical to *R. japonica* infecting humans in China (access. no. CP047359, CP032049) (*7*) *and H. hystricis* (access. no. AP017586 - 8, AP017579) in Japan (*8*).

**Figure.**
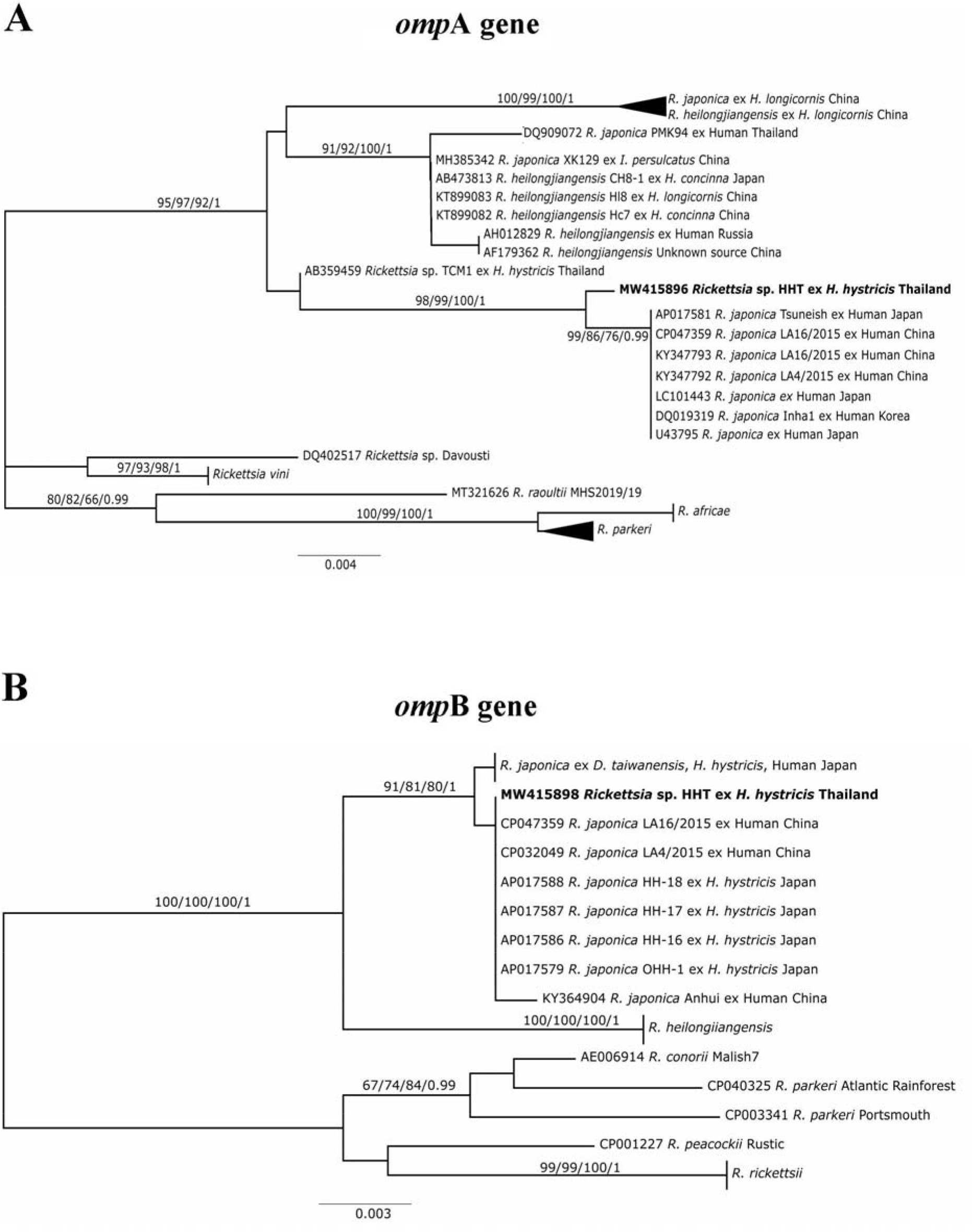
Neighbor-joining trees of *Rickettsia japonica* in *Haemaphysalis hystricis* in Thailand (HHT: indicated in bold characters) based on partial sequences of the *omp*A (A), and *omp*B (B) genes amplified from pooled nymphs of *H. hystricis*. Variability within some clades were collapsed into triangles for visualization. The bootstrap values for neighbor-joining, maximum parsimony, maximum likelihood and probability for Bayesian analysis were shown above branch. Scale bar indicates nucleotide substitutions per site. The name of each sequence containing the GenBank accession number is followed by the name of the *Rickettsia* species, host species and country of origin.

Unfortunately, sequences of *omp*B gene from *H. hystricis* and from humans in Thailand were not available and could not be included in this study. Based on nucleotide BLAST results, the unknown *Rickettsia* sp. detected in the *H. hystricis* ectoparasites of Burmese ferret-badger is most likely *R. japonica*. This is also supported by phylogenetic analyses based on partial *omp*A and *omp*B gene sequences using neighbor-joining, maximum parsimony, maximum likelihood and Bayesian methods. All phylogenetic methods revealed similar tree topologies thus only the neighbor-joining trees are shown (Figure). It is obvious that our *Rickettsia* sp. was clustered with *R. japonica* with strong support (>90%). The results thus clearly demonstrated that the *H. hystricis* nymphs were infected with *R. japonica*. Therefore, the Burmese ferret-badger is a potential reservoir of *R. japonica* which could be transmitted to human via tick vectors such as *H. hystricis* in Thailand.

A variety of tick species are ectoparasites of a wide range of domestic and wild mammals (*9*). The wild fauna may serve as reservoirs of SFG rickettsiae and JSF that can be transmitted to humans through infected ticks (*10*). We have demonstrated for the first time an association of *H. hystricis* ticks infected with *R. japonica* and a wild mammalian host in Thailand. Further investigations on the abundance and distribution of *H. hystricis* ticks parasitizing wild mammals and the prevalence of rickettsial infection in the vectors and hosts are necessary for better understanding the epidemiology of SFG rickettsiae and other tick-borne diseases in Thailand and Southeast Asia.

## Acknowledgments

We thank Dr. Adrian Plant for invaluable comments.

This research was partially supported by Mahidol University, the Thailand Research Fund-Chinese Academy of Science Grant (DBG6180027), and the Center of Excellence on Biodiversity, Office of Higher Education Commission (BDC-PG3-163005 and BDC-PG3-161006).

## About the Author

Dr. Hirunkanokpun is an assistant professor in the Department of Biology, Faculty of Science, Ramkhamhaeng University. Her research interests include ecology of vectors and host-vector-pathogen interactions.

